# ColabSeg: An interactive tool for editing, processing, and visualizing membrane segmentations from cryo-ET data

**DOI:** 10.1101/2023.07.04.547645

**Authors:** Marc Siggel, Rasmus K. Jensen, Julia Mahamid, Jan Kosinski

**Author notes:** Email address (Jan Kosinski).

## Abstract

Cellular cryo-electron tomography (cryo-ET) has emerged as a key method to unravel the spatial and structural complexity of cells in their near-native state at unprecedented molecular resolution. To enable quantitative analysis of the complex shapes and morphologies of lipid membranes, the noisy three-dimensional (3D) volumes must be segmented. Despite recent advances, this task often requires considerable user intervention to curate the resulting segmentations. Here, we present ColabSeg, a Python-based tool for processing, visualizing, cleaning, and fitting membrane segmentations from cryo-ET data for downstream analysis. ColabSeg makes many well-established algorithms for point-cloud processing easily available to the broad community of structural biologists for applications in cryo-ET through its graphical user interface (GUI). We demonstrate the usefulness of the tool with a range of use cases and biological examples. Finally, for a large *Mycoplasma pneumoniae* dataset of 50 tomograms, we show how ColabSeg enables high-throughput membrane segmentation, which can be used as valuable training dataset for fully automated convolutional neural network (CNN)-based segmentation.

## 1. Introduction

In situ cryo-ET has emerged as a powerful method to visualize and analyze the 3D structure of cells and sub-cellular architecture [1, 2, 3, 4]. The 3D volume of a (sub)cellular region, called a tomogram, is reconstructed from 2D projection images acquired on a transmission electron microscope in many different orientations [5, 6]. Macromolecular complexes can be identified in the tomogram, their spatial arrangement can be analyzed in the native environment, and their structure can potentially be determined to near-atomic resolution [1, 2, 3, 4]. Cryo-ET has also proven particularly useful in tracing the abundant cellular membranes in 3D [7, 8]. Membrane segmentations can be used to analyze a variety of membrane properties such as shape, curvature, or volume to understand morphological changes and biological function. These properties can be analyzed with existing software packages [9, 10]. Membrane segmentations can be also exploited to extract associated membrane proteins [11, 12].

However, obtaining these insights comes with numerous challenges. The cryo-ET reconstructions are often noisy or incomplete due to several factors such as (i) the crowded cellular environment, (ii) the missing wedge, (iii) low signal-to-noise ratio, or (iv) artifacts from, i.e., non-vitreous ice or edges of the carbon support film. Due to these complications, segmentation and subsequent analysis of cryo-ET data is still a difficult task and a major bottleneck for fully automated high-throughput analysis of large datasets [4, 5, 13]. Current membrane segmentation methods are still far from fully automated and due to the effort required, many tomograms go unsegmented and are not useful for large-scale statistical analyses.

Various tools and algorithms exist to simplify or partially automate the segmentation of membranes from cryo-ET data. Traditional methods include template matching or a watershed algorithm [14, 15, 16, 17]. A prevalent tool for membrane segmentation, TomoSegMemTV, uses the tensor voting (TV) method [12, 18]. In this six-step pipeline a tomogram is filtered, ridges are enhanced and filtered based on their surfaceness properties. This pipeline and the algorithms are also available as a parallelized implementation for high performance computing (HPC) systems [19]. While the overall performance of the tool is relatively high, it requires tuning many parameters of the pipeline based on the specifics of the tomogram such as the Gaussian filter widths, the size of the neighborhood considered for the tesor voting algorithm, as well as the intensity and surfaceness thresholds. In all cases, the output membrane segmentation often requires laborious manual cleaning with a tool like Amira [20]. More recently, two convolutional neural networks (CNNs) have been developed for particle [21, 22] and membrane segmentation [22]. While these show promise to eventually fully automate the segmentation process, even a state-of-the-art CNN requires large amounts of annotated training data to achieve high-quality segmentations.

To alleviate the above limitations, we developed ColabSeg, an interactive tool for visualizing, editing, and processing membrane segmentations as point clouds from outputs like the tensor voting tool TomoSegMemTV [12] or CNNs [22, 21]. ColabSeg converts the cryo-ET segmentation into point clouds and enables access to a range of well-established computervision algorithms used in processing point clouds such as statistical and eigenvalue-based edge outlier removal [23] without the need for scripting. It also provides tools to fit vesicles and extended membranes for hole-free geometric calculations. The Jupyter notebook in which ColabSeg can be used enables easy remote visualization of the data and is platform-independent. The tool aims to alleviate the tedious and time-consuming task of generating ground truth data for training neural network-based approaches which can drastically improve performance

In this manuscript, we first introduce the general framework, and the software workflow and structure. We explain the features and the foundation of the tools provided for processing and editing segmentation files and showcase the different features of the software on four sample tomograms. Next, we show how ColabSeg can be used to generate a large set of training data for a 3D U-Net DeePiCt [22]. Finally, we discuss possible shortcomings and future directions and give an outlook on how this work might serve as a stepping-stone towards a general CNN for segmentation where no further human intervention is required.

## 2. Methods

### 2.1. Implementation details

ColabSeg is written in Python with accompanying libraries NumPy [24] and SciPy [25]. Features and algorithms of several state-of-the-art point cloud processing libraries, such as Open3D [26] or pyntcloud, are directly used in ColabSeg. The GUI is written using ipywidgets together with py3Dmol [27] to view the point clouds, which can be rendered in any Jupyter notebook. The platform-independent nature of a Jupyter notebook ensures this software package can be run on any cluster or desktop computer with a working Python and Jupyter notebook installation. It can also be installed and used, in principle, remotely on a JupyterHub instance or with a Google Colab notebook. In this way, a notebook can also be run on an external cluster or server and subsequently viewed on a local desktop computer via an ssh connection on-the-go without copying large amounts of data to a local machine. The usage of the tool and its steps are outlined below.

### 2.2. Exemplary workflow for segmenting a reconstructed tomogram

In the following, we describe an exemplary workflow of how to use ColabSeg starting from a reconstructed tomogram (Fig. 1). In step 1, a raw tomogram is processed using existing segmentation software (Fig. 1, step I). This could be a CNN-based segmentation or, in our case, the output from the well-established tensor voting tool TomoSegMemTV [12, 19]. ColabSeg provides a wrapper and GUI for TomoSegMemTV with optimized settings, which proved useful for running TomoSegMemTV for a number of example applications (The exact settings are detailed in the SI). Thus, a reconstructed tomogram is the only required input. All reported settings worked best for binned tomograms with a pixel size of approx. 26 Å, but an acceptable performance was achieved 13 Å as well. We note that a 13-26 Å pixel size is preferred to keep the computational load in an acceptable range. We also provide further instructions for tuning the performance of TomoSegMemTV starting from these optimized settings (see SI). Similar settings appear universally useful, as has been reported by Barad et al. [10] during the preparation of this manuscript. The quality of the overall segmentation and processing with ColabSeg strongly relies on the results of TomoSegMemTV. Therefore optimizing the outputs here has been essential for quality.

**Figure 1:**
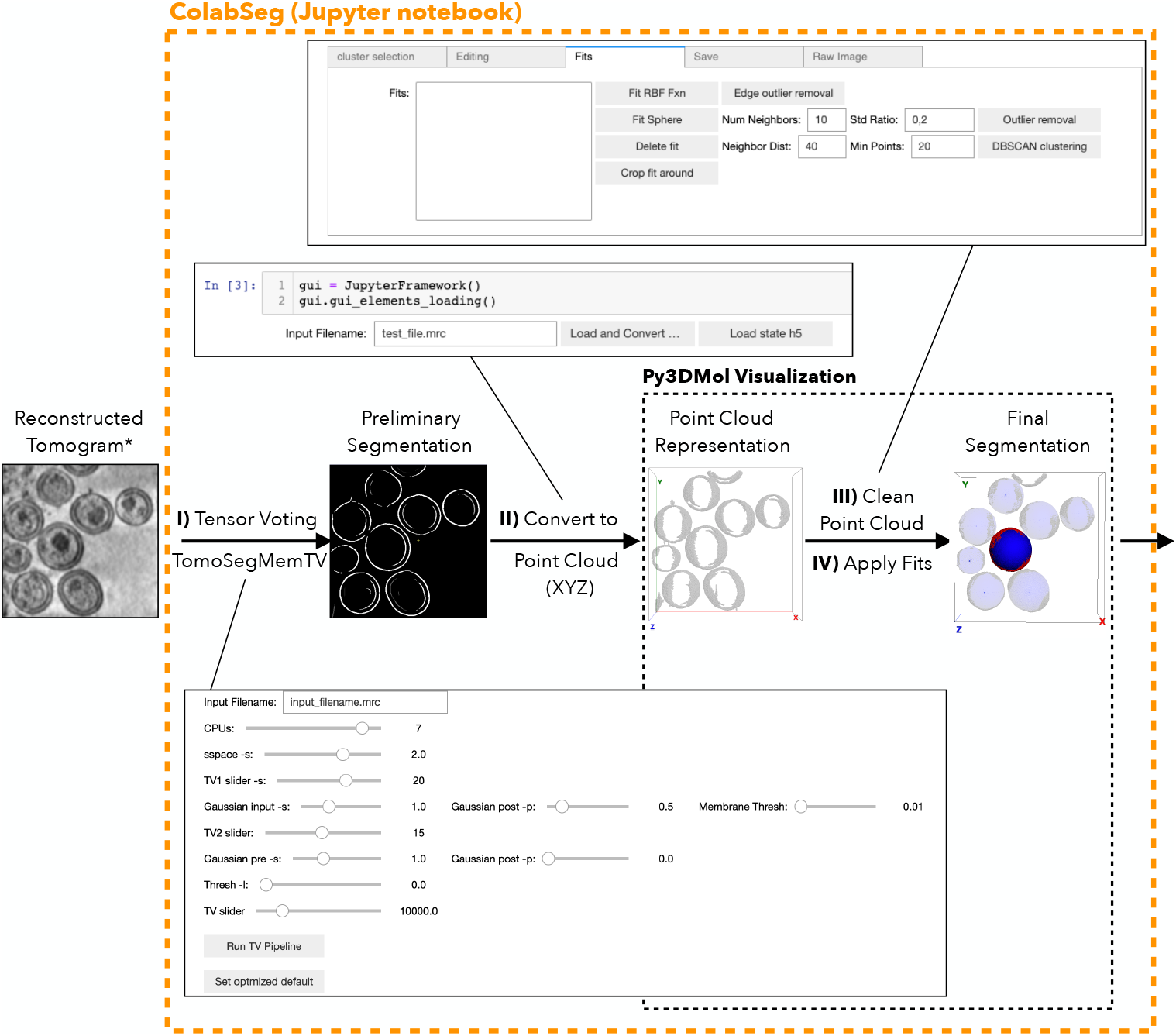
Overview of the Colabseg software and workflow. The demo tomogram contains eight HIV virions (EMDB ID: 13079) [28] and is shown on the left (an image filtered with the Gaussian filter provided in TomoSegMemTV is shown for better visibility). I) The tensor voting workflow [12] is applied to the reconstructed tomogram. This can be accessed and run in the provided GUI. II) The Binary MRC file is then converted to a point cloud file. III) Within the ColabSeg visualizer and editor, the point cloud can be processed, and any artifacts or undesired clusters can be removed using eigenvalue-based edge detection, statistical outlier removal, trimming, and reclustering tools. IV) Fits can be applied to the clusters to interpolate spheres or extended planar membranes to fill holes and extrapolate beyond the data for accurate measurements. The final segmentation can be saved from the main menu as XYZ or (binary) MRC file. Alternatively, a HDF5 state file of the session can be saved throughout the editing process to resume editing later.

We find that permissive thresholds in both the membrane detection and thresholding step in the settings (see SI) are advantageous to achieve more detailed and complete segmentations, even of smaller membrane features. While in this case more noise and false positives are often picked up, many of these excess segments can be removed using the functions of ColabSeg at a later stage. Manual and semi-automated removal of artifacts or false-positive is considerably faster than post-segmentation manual addition of missing features.

In step 2, ColabSeg takes as input a binary segmentation MRC file generated from TomoSegMemTV, where voxels are labeled with integer values corresponding to their cluster. Voxels, where no membrane is present, are labeled zero and disregarded. ColabSeg converts these clusters from voxels to a cloud of XYZ point coordinates. Each of the integer values tagged voxel is assigned to a separate point cloud in accordance with the clustering output from the connected-component algorithm provided in TomoSegMemTV (Fig. 1, Step II). This input is the default output of the cluster analysis from TomoSegMemTV. Inputs from other software need to be prepared similarly but, at minimum, be a binary file. Alternatively, the DBSCAN re-clustering algorithm included in ColabSeg can be used to assign initial clusters as well, albeit with a result different from the connected component algorithm used by TomoSegMemTV.

The point clouds and clusters can be directly visualized in the notebook using an interactive py3Dmol viewer [27]. Point cloud coordinates are imported as dummy atoms. Importantly, to avoid memory issues due to a large number of points, the point cloud density is downsampled for visualization purposes. The degree of downsampling is adjusted automatically depending on the overall count of points in the point cloud.

In step 3, the point cloud derived from the segmentation can be cleaned and further refined using several utilities (Fig. 1, Step III). Detailed usage is described below and shown with different examples. This entails rapidly merging and deleting clusters, which is also possible in Amira, but avoids having to switch the software throughout the workflow and requiring another desktop software [20]. Clusters can be further split using the DBSCAN clustering algorithm. Point clouds can be filtered using a statistical outlier method or eigenvalue-based edge detection. The edges of the tomograms can be trimmed to reduce noise at the fringes of the tomograms. All methods can be combined to achieve a high-quality segmentation.

Finally, in step 4, idealized planar and spherical shapes can be fitted to planar membranes and spherical viruses or vesicles (Fig. 1, step IV). These two methods allow the fitting of a broad range of structures in biological samples and can be used to fill holes in existing segmentations, as required for quantitative spatial analysis. The details and usage are described below with the real biological sample data.

ColabSeg also allows saving the processed segmentations in various formats. Most commonly, MRC files can be written for use in further analysis pipelines. Additionally, the segmentations can be written as XYZ points or text files for subsequent custom analysis or visualization in other software packages such as VMD [29]. It is also possible to save and load custom ColabSeg state files in HDF5 file format at any time during processing to save intermediate steps. All selections and settings of the GUI are saved and can be reconstructed from this file.

### 2.3. Advanced features for editing and cleaning tomograms

ColabSeg provides access to numerous point cloud processing algorithms commonly used in computer graphics to refine and correct false positives and split clusters. To demonstrate a possible use of these algorithms we prepared a synthetic demo dataset consisting of a point cloud with two intersecting planes, representative of membranes, and Gaussian noise added to some of the points (Fig. 2) to demonstrate the features. TomoSegMemTV [12] and other tools often include numerous false positives outside the cellular region, such as ice contaminants or fiducial markers. Additionally, at the edge of the tomogram or cellular slice (in the case of focused ion beam lamellae), the data is ambiguous, resulting in some noise. Moreover, point clouds are occasionally falsely grouped by the connected component algorithm provided by TomoSegMemTV [12] due to noise.

**Figure 2:**
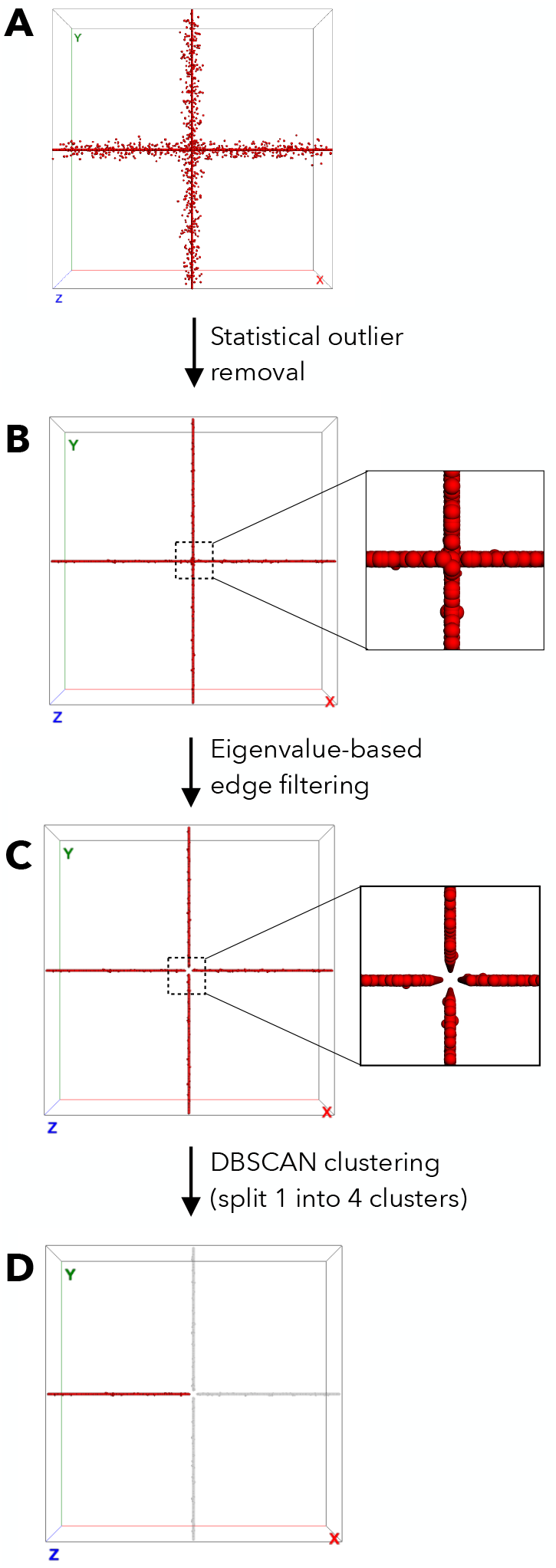
Overview of implemented point cloud processing and cleaning features using a synthetic MRC file. Shown are snapshots from the ColabSeg 3D viewer. A selected cluster is shown in red. (A) The initial file consists of two intersecting planes with 10000 points having Gaussian noise added to them. The initial synthetic MRC file has a cross shape. (B) Statistical outlier removal for data cleaning as implemented in the Open3D library. (C) Eigenvalue-based edge filtering example. (D7) DBSCAN algorithm for cluster separation. All features can be combined in various order to solve more complex segmentation issues resulting from inaccuracies in the tensor voting. The view render in ColabSeg dynamically downsamples the number of points to avoid speed and memory issues.

To remedy such issues, we provide statistical outlier removal based on the implementation in Open3D [26] (Fig. 2). The free parameters include the number of the k-nearest neighbors, which are used to calculate the average distance for a given point. The threshold is given by the standard deviation from the distribution of distance, where lower values result in stricter cutoffs. Points that constitute background noise are removed.

We also provide a more advanced eigenvalue-based outlier removal methodology. Bazazian et al. [23] show that the eigenvalues of the covariance matrix that are defined by each point’s k-nearest neighbors are an effective way of filtering points constituting an edge with a sharp change in curvature. For most examples in biological systems which are flat, extended membranes such as plasma membranes, nuclear envelopes, and other large structures, this method works particularly well (see below Fig. 4C). Since membranes are most commonly represented in smooth extended surfaces, most artifacts manifest as unordered, highly distorted point clouds. This analysis outperforms other methods for edge detection such as PCA analysis or triangulation-based methods combined with clustering, as reported in ref. [23]. It does not rely on the initial clustering step, which would add additional free parameters to the analysis and depend highly on the point cloud density. In our test case (Fig 2) the intersection of the two planes is easily filtered with this approach. Good results are also observed for spherical vesicles or tubular structures if large enough continuous pieces of the membrane are captured in the reconstructed volume (Fig 4).

To enable the re-clustering of clusters initially assigned through the connected-component algorithm in TomoSegMemTV, we provide the DB-SCAN algorithm [30] which is implemented in the open3D library [26]. In our test case (Fig. 2) the initial single cluster can now be split into four separate clusters. This approach was also chosen for some of the applications shown below (Fig. 4B). In real applications, tuning of the parameters also enables the filtering of outlier points [30].

TomoSegMemTV [12] and other tools often include numerous false positives outside of the cellular region. These can easily be removed by trimming the edge using a corresponding tool in ColabSeg (Supporting Information Fig. S4). Removing points at the respective upper and lower bound drastically improves the quality of the segmentation. Finally, in case the cellular or lamellae regions are not rotation-corrected at reconstruction step, this can be done in ColabSeg by performing a plane fit through all points of the membrane and subsequently aligning with the z-axis (SI Fig. S4).

### 2.4. Fitting procedures for hole filling

Many analyses of cryo-ET data rely on spatial analysis such as Euclidian distances, e.g., from nearest membranes. Here complete, hole-free membrane representations are required to achieve correct and robust results. However, many segmentations have holes or missing pieces which impede these calculations, which is another reason for many researchers to resort to manual segmentations. Therefore, we provide access to radial basis function (RBF) plane fits for extended membranes and sphere fits for vesicles (Fig. 3) - two common geometries for membrane structures found in tomograms. These enable extrapolating and filling missing pieces in segmentations for accurate calculations. Within the GUI, one or multiple segments can be selected, and the fit is automatically performed. The SciPy [25] RBF fit and ‘lstsq’ function are used to perform the fits.

**Figure 3:**
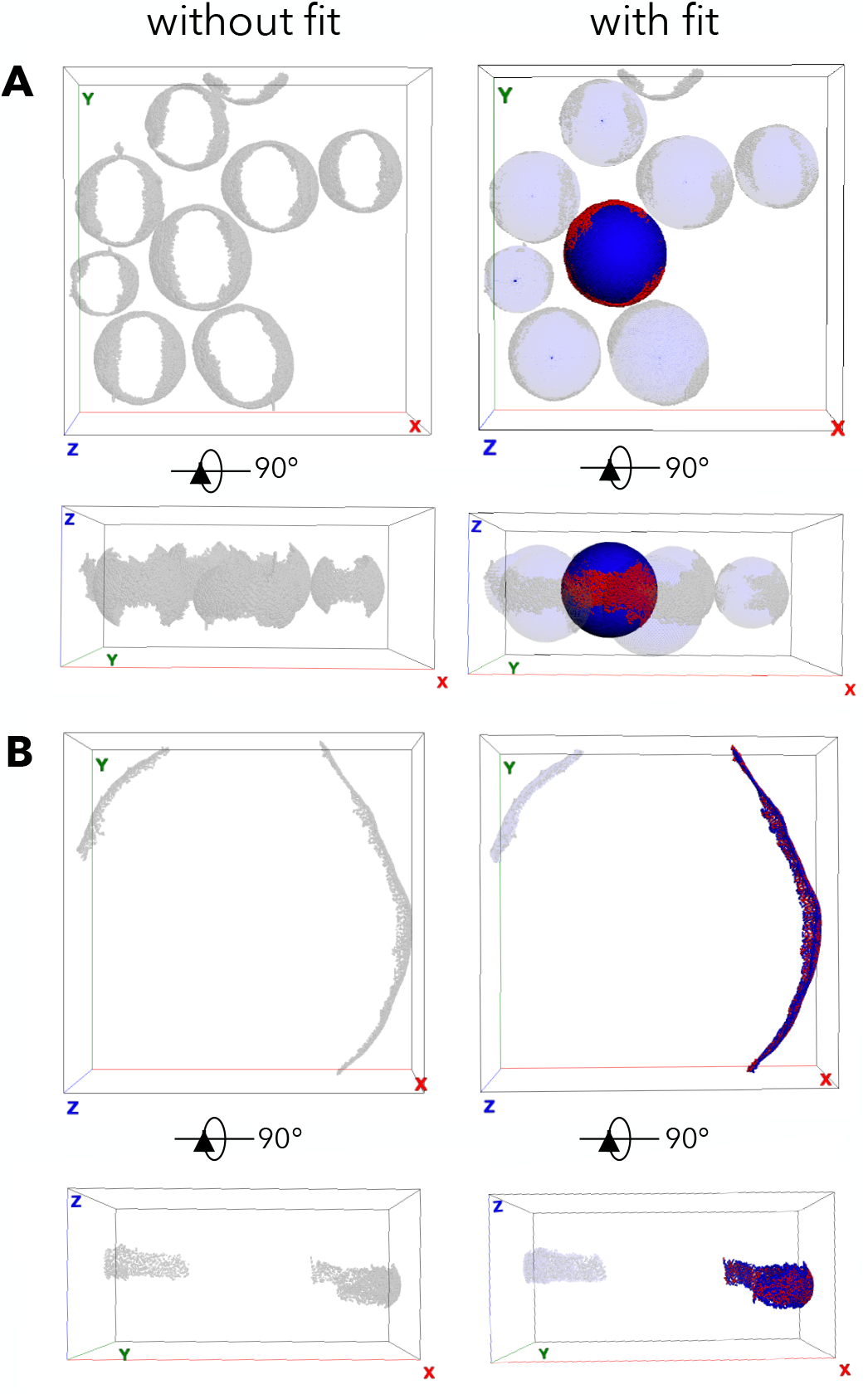
Examples of sphere and radial basis function fit functionality in ColabSeg. (A) Sphere fits for eight HIV virions. The fit is shown in (light) blue. The original segmentation derived from the cryo-ET data is shown in grey. An exemplary virion cluster and the corresponding fit are selected in dark red and blue, respectively. A top and side view snapshots from the ColabSeg viewer are shown before and after the processing. (B) Radial basis function fit on two outer membranes of *Mycoplasma pneumoniae*. The initial segmentation is shown in grey, and the fits in blue. One exemplary cluster and corresponding fit are shown in dark red and blue. The fit was trimmed to only include points within 50 Å of the input data. A top and side view snapshot from the ColabSeg viewer is shown before and after the processing.

### 2.5. Using ColabSeg to generate training data for the CNN of DeePiCt

The membranes of ten *Mycoplasma pneumoniae* tomograms were segmented manually using Amira (version 2022.1, Thermo Fischer). In addition, membranes in 50 tomograms were segmented using TomoSegMemTV [12] and cleaned using Colabseg (detailed below). The segmentations were provided as input for training the DeePiCt model [22], using either five manually segmented tomograms from Amira, as well as 5, 10, 25, or 50 of the semi-automatically segmented tomograms from ColabSeg. The remaining five manually segmented tomograms were used for validation. The DeePiCt model was trained at pixel size 13.6 Å/pixel with box size and overlap of 64 and 6 pixels, respectively. The default U-net convolutional network of DeePiCt [22] was used, with a depth of 2 using 16 initial convolutions. In each epoch, 80% of the data was used for training and validated using the dice coefficient against the remaining 20% of the data. A total of 300 epochs were run for training each model, but training normally converged within the first 50 epochs. The resulting models were used to predict the membrane in the validation set, and the output was filtered to a binarized map using a threshold of 0.4, excluding any cluster containing less than 1000 voxels.

## 3. Results and Discussion

### 3.1. Biological examples for editing and fitting segmentations

We tested ColabSeg to refine membrane segmentations from various cryo-ET datasets. Our examples span HIV virions [28], the inner membrane complex (IMC) of Plasmodium falciparum [31], and *Mycoplasma pneumoniae* cells. Additionally, an early version of the tool has been used to segment plasma membranes [32]. Starting from reconstructed tomograms, we use ColabSeg together with TomoSegMemTV [12] to arrive at fitted target segmentations (Fig. 4) For the commonly used HIV virions dataset (EMDB ID: 13079, [28]), we first used TomoSegMemTV to segment viral membranes with a low threshold (Fig 4A). In some instances, two virion membranes were determined as one cluster by the TomoSegMemTV connected component algorithm. Therefore, filtering using the statistical outlier tool and reclustering with the DBSCAN algorithm was necessary. We also used ColabSeg to remove clusters that are not virion membranes. Subsequently, we applied sphere fit to all virions. These could enable further distance measurements and the use of the fit parameters for spherically constrained 3D subtomogram averaging [33].

**Figure 4:**
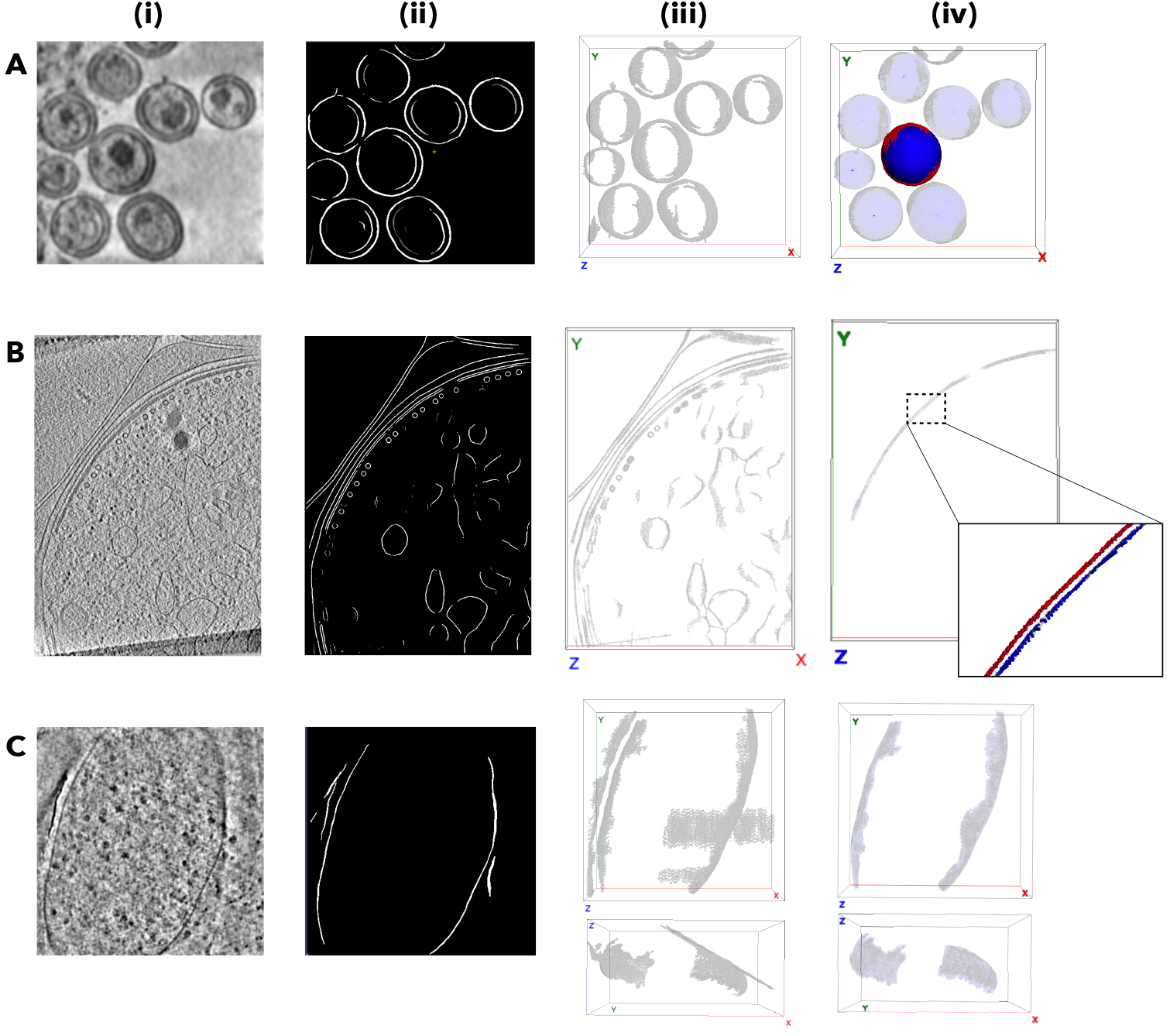
Application of ColabSeg on biological example systems. The figures show: (A) the separation and fit of HIV virions (EMDB ID: 13079, [28], (B) the extraction of Plasmodium falciparum inner membrane complex (IMC) for further analysis as used in Ferreira, Prazak et al. [31], and, (C) the clean-up and artefact removal from a *Mycoplasma pneumoniae* tomogram. All Figures show (from left to right) a central slice from the reconstructed tomogram (shown is the filtered first step of the TomoSegMemTV pipeline) (i), the output of the TomoSegMemTV tool (ii), the imported segmentation in ColabSeg visualized as point cloud (iii), and the final processed and fit point cloud for further use in analysis (iv). Here, grey points indicate the processed data, blue the fits, and red highlights a cluster of interest. Each sample data set combines a number of features including cluster merging, deletion, edge filtering, statistical outlet removal and various fits. The detailed workflow for each of the images is outlined in the provided user guide.

Next, we used the tools from ColabSeg to extract the inner membrane complex (IMC) of Plasmodium falciparum to enable accurate measurements of their distance to microtubules (Fig. 4B) [32]. Here, we first ran TomoSeg-MemTV to extract the main features (plasma and intracellular membranes, and microtubules), and then used a combination of edge-filtering, edge trimming, statistical outlier removal, and DBSCAN re-clustering to select and segment both bilayers of the IMC [31]. Workflows using solely TomoSeg-MemTV resulted in either patchy membranes or large amounts of excess noise which would have interfered with the measurement (Fig. 4B). In particular, tomograms from thin lamella profited from the edge trimming procedure, in most cases removing a bulk of false positive noise. We then used the RBF fit function to fill any holes in the membranes to avoid errors in the distance measurements. These measurements were then used to show the high degree of consistency in membrane-microtubule distances, indicating the presence of a linker, which would not have been possible without accurate segmentation of the IMC [31].

Finally, we demonstrate the utility of the eigenvalue-based edge filtering on problematic tomograms from a data set of *Mycoplasma pneumoniae* (Fig. 4C). Sometimes, edges of the grid support film or other high contrast features, are often falsely picked up in the segmentation process resulting in unreasonable intersections (Fig. 4C, left (i, ii)). When using TomoSegMemTV, this leads to artifacts that so far have had to be separated manually. Using the eigenvalue edge filtering we can remove such artifacts from the segmentations (Fig. 4C, right (iii, iv)): the edge detection removed points along the distinct edge where the feature intersects with the membrane. Subsequently, we were able to split these two clusters by using the DBSCAN clustering algorithm and using the RBF fit function to fill the resulting hole (Fig. 4C, right).

These examples demonstrate the utility and ease of use of ColabSeg for a variety of tasks. By combining different features of the software, we were able to resolve a broad range of issues resulting from the initial segmentations and could enable accurate analysis of many tomograms in a short amount of time. Overall, we note that ColabSeg is strongly dependent on the initial segmentation. The quality of the segmentations, by default, the output from TomoSegMemTV [12], is a decisive factor for the successful segmentation of membranes. For this reason, we tested various settings for TomoSegMemTV and provide the best ones to the user. We found TomoSegMemTV performs particularly well with a permissive threshold in the membrane ‘surfaceness’ step to capture all membrane features irrespective of their size. Additionally, thick cellular regions or lamellae have a particularly low signal-to-noise ratio and usually result in poorer segmentations which are difficult to clean. In most cases, some manual intervention will be required to clean the remaining segmented clusters, and ColabSeg greatly facilitates this task.

### 3.2. Training data generated by ColabSeg improve deep learning performance

ColabSeg was developed not only to ease the semi-automated segmentation and directly enable analysis of the data, but also to generate data for training convolutional neural network (CNN) approaches faster and increasing throughput of cryo-ET data analysis. Recently, open-source software solutions for the segmentation of tomograms using 2D and 3D U-Nets have been made available [22, 21]. The quality of the segmentations from the CNNs strongly depends on the available training data. To date, most CNNs designed for cryo-ET rely on very few (one to ten) segmented tomograms as training data, which are often generated manually with software like Amira. The resulting trained models then only perform well for a narrow scope of similar data, if at all. To achieve larger transferability and higher quality segmentation we propose using the ColabSeg pipeline to quickly generate a sufficiently large training set for CNN-based approaches.

To test this, we used the ColabSeg pipeline in conjunction with the recently developed DeePiCt network [22]. First, we use ColabSeg to segment several hundreds of tomograms from *Mycoplasma pneumoniae*, filtering artifacts and false positive cases with the methods introduced above. By doing so, we prepared several hundred segmented and cleaned tomograms for training data. Next, we trained DeePiCt on several different subsets of the clean data as well as some manual segmentations described above to assess differences in performance. A set of 5 manually annotated tomograms were used as a validation data set.

We found that using a large number of tomograms for training drastically improves the performance of the network for the same validation dataset (Fig. 5). Importantly, a dataset consisting of tomograms solely curated with ColabSeg outperformed a data set segmented manually using Amira. With this larger training data set, we were able to reliably filter out artifactual segmentation of the support film edge, which is visible in many tomograms. The corresponding DeePiCt model could reliably segment membranes from unseen data from the *mycoplasma* data set at a level similar to manual annotation.

**Figure 5:**
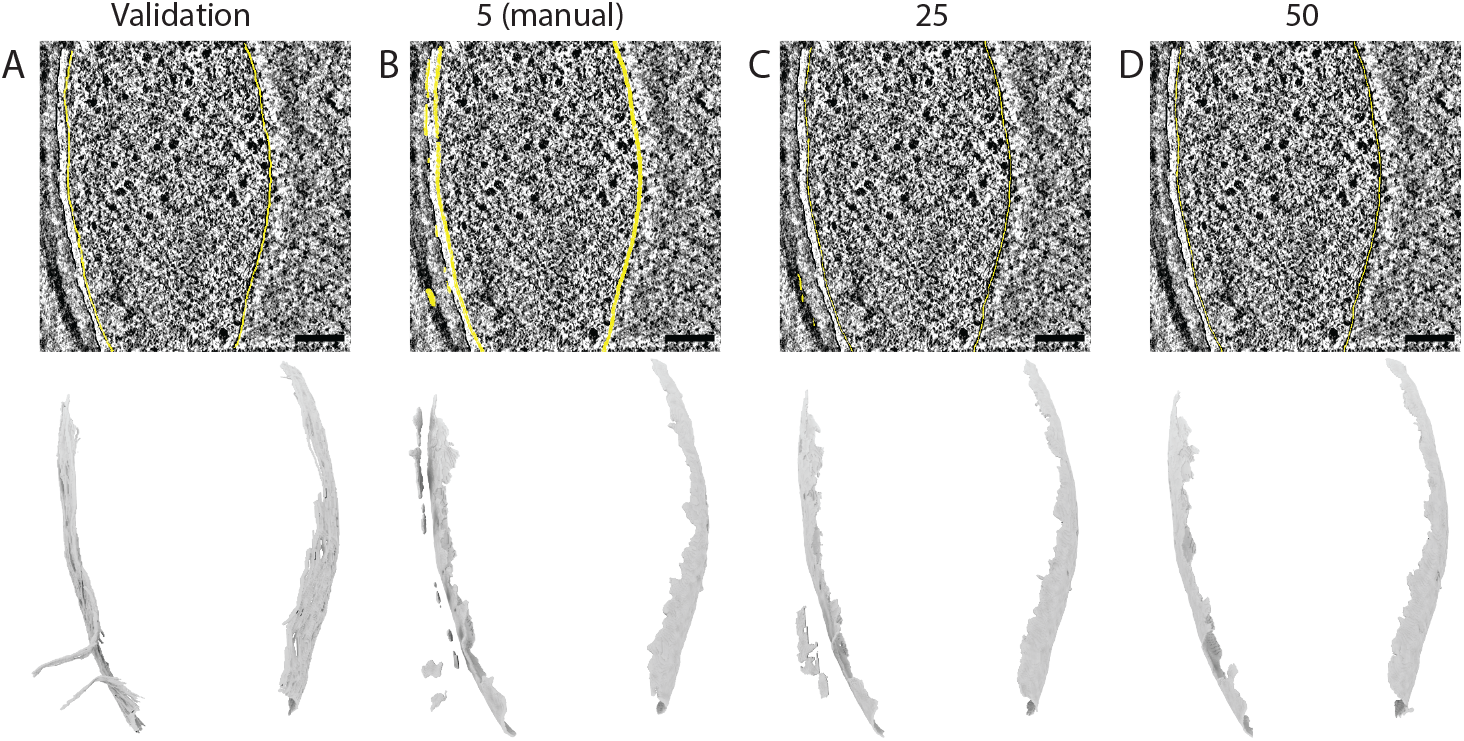
Use of ColabSeg data to improve CNN training for fully automated membrane segmentation. The top row shows a 2D slice of the tomogram of *Mycoplasma pneumoniae* with the segmentation as yellow overlay. The bottom row shows 3D volume render of the entire segmentation. (A) Slice of the tomogram deconvolved with Warp from the validation data set for machine learning. The ground truth was generated by human manual annotation. (B) Results for a CNN trained on 5 manually annotated tomograms. (C and D) Results for the same CNN trained on 25 (C) or 50 (D) tomograms segmented and processed with the ColabSeg tool.

*Mycoplasma pneumoniae* is a comparably straightforward test system. Since it is a small bacterium and contains no complex membrane-bound organelles, the segmentation task is significantly easier than for other cell types. Similarly, viruses such as HIV, influenza A virus, Ebola, or other simple membrane-bound structures can easily be analyzed using ColabSeg. Eukaryotic cells pose a much bigger challenge for accurate segmentation, for example, the highly curved membranes such as the tubule or the cristae of mitochondria. Here, a combination of computer-aided and manual segmentation in ColabSeg will enable gathering training data of sufficient quality for accurate CNN-based segmentations of such cellular structures.

## 4. Conclusions and Outlook

We presented ColabSeg, a Jupyter notebook-based GUI for visualizing, editing, and post-processing segmentations of membranes in cryo-ET data. We enable users to generate initial segmentations with TomoSegTV implemented in our tool, quickly filter relevant segments, remove noise as well as generate fits to fill holes, and to enable downstream quantitative measurements with other tools. We provide numerous algorithms used in point cloud processing accessible in a simple interface without requiring any scripting. Advanced users can build semi-automatic workflows by using the provided clases and the accompanying image-processing libraries. We show that even though some semi- and fully automatic methods for segmentation exist, user intervention and curation are still necessary to arrive at high-quality segmentations. We demonstrated the usefulness of ColabSeg on a range of use cases, which are being used to address a broad range of scientific questions [31, 32]. Finally, we show how ColabSeg can be used in conjunction with a CNN, DeePiCt, for generating sufficient training data for more accurate automated segmentation.

Currently, ColabSeg is designed and set up to visualize and process only membrane segmentations. However, the GUI can be extended in a straight-forward way to include visualization of other biomolecules such as proteins, their densities, and even the accompanying structural models. Since the underlying 3D visualization software, py3Dmol, was intended to represent proteins and can handle complex representation [27], it might also become a useful tool to visualize and analyze protein complexes, measure their arrangement, and analyze their properties in segmented cryo-ET densities.

The backend functionality can easily be adapted to interface with other visualizing software packages with some minor changes to the interface. It could be included in existing image processing such as the Napari viewer [34] for broader applicability and to give access to more features in the future. The lightweight design of the software and browser-based visualization possibilities might be deployed on web servers or on Google Colab for easy access and analysis. This might make sharing these algorithms, developing and deploying online servers for the processing of data easier.

In the future, ColabSeg in conjunction with other segmentation methods will hopefully speed up the process of membrane segmentation and as a result, help generate a larger annotated set of cryo-ET data. We envision this tool as a stepping stone to lower the barrier for experimentalists to annotate their raw tomograms and thus populate a growing body of data for use as training data for deep learning methods such as DeePiCt or DeepFinder [22, 21], which is still lacking to date. Eventually, CNN models will be trained on sufficiently broad data such that near error-free and fully automated segmentation should be available in the future.

## 5. Software Availability

ColabSeg is available under an Apache License 2.0 license from GitHub (https://github.com/KosinskiLab/colabseg) free of charge and can be installed as a Python package. It can be used from a Jupyter notebook or Colab notebook (setup required by the user). Third-party contributions are welcome as pull requests on GitHub. Further analysis or processing features can be easily added to the existing GUI. Developers can add functions in the backend and add additional GUI widgets to the code to access their functions. Methods can use one or multiple clusters of points as input.

## Supporting information

User Guide

## 6. Acknowledgements

The work was supported by EMBL and a research fellowship from the EMBL Interdisciplinary Postdoc (EIPOD) Programme under Marie Curie Cofund Actions MSCA-COFUND-FP (grant agreement number: 847543) to MS. RKJ was supported by a postdoctoral fellowship from the DFF (grant number 0164-00010A). We thank Dr. Josie L. Ferreira for providing multiple tomograms of P. Falciparum stages and for helpful discussions. We thank Liang Xue for providing multiple tomograms of *Mycoplasma pneumoniae*. We thank Herman Fung, Veijo T. Salo, Sara K. Goetz, and Anastasiia Babenko for testing the software and critical feedback. We thank Thomas Hofmann for discussions and together with the rest of EMBL IT providing the IT infrastructure.

## References

[1] J. Mahamid, S. Pfeffer, M. Schaffer, E. Villa, R. Danev, L. Kuhn Cuellar, F. Forster, A. A. Hyman, J. M. Plitzko, W. Baumeister, Visualizing the molecular sociology at the HeLa cell nuclear periphery, Science (80-.). 351 (6276) (2016) 969–972. doi:10.1126/science.aad8857.

[2] S. Pfeffer, J. Dudek, M. Schaffer, B. G. Ng, S. Albert, J. M. Plitzko, W. Baumeister, R. Zimmermann, H. H. Freeze, B. D. Engel, F. Förster, Dissecting the molecular organization of the translocon-associated protein complex, Nat. Commun. 8 (1) (2017) 14516. doi:10.1038/ncomms14516.

[3] F. Wilfling, C.-W. Lee, P. S. Erdmann, Y. Zheng, D. Sherpa, S. Jentsch, B. Pfander, B. A. Schulman, W. Baumeister, A Selective Autophagy Pathway for Phase-Separated Endocytic Protein Deposits, Mol. Cell 80 (5) (2020) 764–778.e7. doi:10.1016/j.molcel.2020.10.030.

[4] V. Lučić, A. Rigort, W. Baumeister, Cryo-electron tomography: The challenge of doing structural biology in situ, J. Cell Biol. 202 (3) (2013) 407–419. doi:10.1083/jcb.201304193.

[5] E. Pyle, G. Zanetti, Current data processing strategies for cryo-electron tomography and subtomogram averaging, Biochem. J. 478 (10) (2021) 1827–1845. doi:10.1042/BCJ20200715.

[6] N. Volkmann, Methods for Segmentation and Interpretation of Electron Tomographic Reconstructions, in: Methods Enzymol., 1st Edition, Vol. 483, Elsevier Inc., 2010, pp. 31–46. doi:10.1016/S0076-6879(10)83002-2.

[7] W. Wietrzynski, M. Schaffer, D. Tegunov, S. Albert, A. Kanazawa, J. M. Plitzko, W. Baumeister, B. D. Engel, Charting the native architecture of chlamydomonas thylakoid membranes with single-molecule precision, Elife (2020). doi:10.7554/eLife.53740.

[8] D. Zabeo, K. M. Davies, Studying membrane modulation mechanisms by electron cryo-tomography, Curr. Opin. Struct. Biol. 77 (2022) 102464. doi:10.1016/j.sbi.2022.102464.

[9] M. Salfer, J. F. Collado, W. Baumeister, R. FernándezBusnadiego, A. Martínez-Sánchez, Reliable estimation of membrane curvature for cryo-electron tomography, PLoS Comput. Biol. 16 (8) (2020) 1–29. doi:10.1371/journal.pcbi.1007962.

[10] B. A. Barad, M. Medina, D. Fuentes, R. L. Wiseman, D. A. Grotjahn, Quantifying organellar ultrastructure in cryo-electron tomography using a surface morphometrics pipeline, J. Cell Biol. 222 (4) (apr 2023). doi:10.1083/jcb.202204093.

[11] L. Lamm, R. D. Righetto, W. Wietrzynski, M. Pöge, A. Martinez-Sanchez, T. Peng, B. D. Engel, MemBrain: A deep learning-aided pipeline for detection of membrane proteins in Cryo-electron tomograms, Comput. Methods Programs Biomed. 224 (2022) 106990. doi:10.1016/j.cmpb.2022.106990.

[12] A. Martinez-Sanchez, I. Garcia, S. Asano, V. Lucic, J.-J. Fernandez, Robust membrane detection based on tensor voting for electron tomography, J. Struct. Biol. 186 (1) (2014) 49–61. doi:10.1016/j.jsb.2014.02.015.

[13] X. Wu, X. Zeng, Z. Zhu, X. Gao, M. Xu, Template-Based and Template-Free Approaches in Cellular Cryo-Electron Tomography Structural Pattern Mining, in: Comput. Biol., Codon Publications, 2019, pp. 175–186. doi:10.15586/computationalbiology.2019.ch11.

[14] M. N. Lebbink, W. J. Geerts, T. P. van der Krift, M. Bouwhuis, L. O. Hertzberger, A. J. Verkleij, A. J. Koster, Template matching as a tool for annotation of tomograms of stained biological structures, J. Struct. Biol. 158 (3) (2007) 327–335. doi:10.1016/j.jsb.2006.12.001.

[15] M. N. Lebbink, W. J. Geerts, E. van Donselaar, B. M. Humbel, J. A. Post, L. O. Hertzberger, A. J. Koster, A. J. Verkleij, Electron tomography and template matching of biological membranes, in: EMC 2008 14th Eur. Microsc. Congr. 1-5 Sept. 2008, Aachen, Ger., Springer Berlin Heidelberg, Berlin, Heidelberg, 2008, pp. 83–84.

[16] S. F. Tasel, E. U. Mumcuoglu, R. Z. Hassanpour, G. Perkins, A validated active contour method driven by parabolic arc model for detection and segmentation of mitochondria, J. Struct. Biol. 194 (3) (2016) 253–271. doi:10.1016/j.jsb.2016.03.002.

[17] I. Luengo, M. C. Darrow, M. C. Spink, Y. Sun, W. Dai, C. Y. He, W. Chiu, T. Pridmore, A. W. Ashton, E. M. Duke, M. Basham, A. P. French, SuRVoS: Super-Region Volume Segmentation workbench, J. Struct. Biol. 198 (1) (2017) 43–53. doi:10.1016/j.jsb.2017.02.007.

[18] Wai-Shun Tong, Chi-Keung Tang, P. Mordohai, G. Medioni, First order augmentation to tensor voting for boundary inference and multiscale analysis in 3d, IEEE Trans. Pattern Anal. Mach. Intell. 26 (5) (2004) 594–611. doi:10.1109/TPAMI.2004.1273934.

[19] J. J. Moreno, E. M. Garzon, J. J. Fernández, A. Martínez-Sánchez, HPC enables efficient 3D membrane segmentation in electron tomography, J. Supercomput. (2022). doi:10.1007/s11227-022-04607-z.

[20] D. Stalling, M. Westerhoff, H. C. Hege, Amira: A highly interactive system for visual data analysis, in: Vis. Handb., 2005. doi:10.1016/B978-012387582-2/50040-X.

[21] E. Moebel, A. Martinez-Sanchez, L. Lamm, R. D. Righetto, W. Wietrzynski, S. Albert, D. Lariviere, E. Fourmentin, S. Pfeffer, J. Ortiz, W. Baumeister, T. Peng, B. D. Engel, C. Kervrann, Deep learning improves macromolecule identification in 3D cellular cryo-electron tomograms, Nat. Methods 18 (11) 2021) 1386–1394. doi:10.1038/s41592-021-01275-4.

[22] I. de Teresa-Trueba, S. K. Goetz, A. Mattausch, F. Stojanovska, C. E. Zimmerli, M. Toro-Nahuelpan, D. W. C. Cheng, F. Tollervey, C. Pape, M. Beck,A. Diz-Munõz, A. Kreshuk, J. Mahamid, J. B. Zaugg, Convolutional networks for supervised mining of molecular patterns within cellular context, Nat. Methods 20 (2) (2023) 284–294. doi:10.1038/s41592-022-01746-2.

[23] D. Bazazian, J. R. Casas, J. Ruiz-Hidalgo, Fast and Robust Edge Extraction in Unorganized Point Clouds, in: 2015 Int. Conf. Digit. Image Comput. Tech. Appl., IEEE, 2015, pp. 1–8. doi:10.1109/DICTA.2015.7371262.

[24] C. R. Harris, K. J. Millman, S. J. van der Walt, R. Gommers, P. Virtanen, D. Cournapeau, E. Wieser, J. Taylor, S. Berg, N. J. Smith, R. Kern, M. Picus, S. Hoyer, M. H. van Kerkwijk, M. Brett, A. Haldane, J. F. del Río, M. Wiebe, P. Peterson, P. Gerard-Marchant, K. Sheppard, T. Reddy, W. Weckesser, H. Abbasi, C. Gohlke, T. E. Oliphant, Array programming with NumPy, Nature 585 (7825) (2020) 357–362. arXiv:2006.10256, doi:10.1038/s41586-020-2649-2.

[25] P. Virtanen, R. Gommers, T. E. Oliphant, M. Haberland, T. Reddy, D. Cournapeau, E. Burovski, P. Peterson, W. Weckesser, J. Bright, S. J. van der Walt, M. Brett, J. Wilson, K. J. Millman, N. Mayorov, A. R. J. Nelson, E. Jones, R. Kern, E. Larson, C. J. Carey, I. Polat, Y. Feng, E. W. Moore, J. VanderPlas, D. Laxalde, J. Perktold, R. Cimrman, I. Henriksen, E. A. Quintero, C. R. Harris, A. M. Archibald, A. H. Ribeiro, F. Pedregosa, P. van Mulbregt, A. Vijaykumar, A. P. Bardelli, A. Rothberg, A. Hilboll, A. Kloeckner, A. Scopatz, A. Lee, A. Rokem, N. Woods, C. Fulton, C. Masson, C. Häggström, C. Fitzgerald, A. Nicholson, D. R. Hagen, D. V. Pasechnik, E. Olivetti, E. Martin Wieser, F. Silva, F. Lenders, F. Wilhelm, G. Young, G. A. Price, G.-L. Ingold, G. E. Allen, G. R. Lee, H. Audren, I. Probst, J. P. Dietrich, J. Silterra, J. T. Webber, J. Slavič, J. Nothman, J. Buchner, J. Kulick, J. L. Schönberger, J. V. de Miranda Cardoso, J. Reimer, J. Harrington, J. L. C. Rodríguez, J. Nunez-Iglesias, J. Kuczynski, K. Tritz, M. Thoma, M. Newville, M. Kümmerer, M. Bolingbroke, M. Tartre, M. Pak, N. J. Smith, N. Nowaczyk, N. Shebanov, O. Pavlyk, P. A. Brodtkorb, P. Lee, R. T. McGibbon, R. Feldbauer, S. Lewis, S. Tygier, S. Sievert, S. Vigna, S. Peterson, S. More, T. Pudlik, T. Oshima, T. J. Pingel, T. P. Robitaille, T. Spura, T. R. Jones, T. Cera, T. Leslie, T. Zito, T. Krauss, U. Upadhyay, Y. O. Halchenko, Y. Vázquez-Baeza, SciPy 1.0: fundamental algorithms for scientific computing in Python, Nat. Methods 17 (3) (2020) 261–272. arXiv:1907.10121, doi:10.1038/s41592-019-0686-2.

[26] Q.-Y. Zhou, J. Park, V. Koltun, Open3D: A Modern Library for 3D Data Processing (jan 2018). arXiv:1801.09847.

[27] N. Rego, D. Koes, 3Dmol.js: Molecular visualization with We-bGL, Bioinformatics 31 (8) (2015) 1322–1324. doi:10.1093/bioinformatics/btu829.

[28] S. Mattei, A. Tan, B. Glass, B. Müller, H. G. Kräusslich, J. A. Briggs, High-resolution structures of HIV-1 Gag cleavage mutants determine structural switch for virus maturation, Proc. Natl. Acad. Sci. U. S. A. 115 (40) (2018) E9401–E9410. doi:10.1073/pnas.1811237115.

[29] W. Humphrey, A. Dalke, K. Schulten, VMD: Visual molecular dynamics, J. Mol. Graph. 14 (1) (1996) 33–38. doi:10.1016/0263-7855(96)00018-5.

[30] M. Ester, H.-P. Kriegel, J. Sander, X. Xu, A Density-Based Algorithm for Discovering Clusters in Large Spatial Databases with Noise, in: KDD, 1996.

[31] J. L. Ferreira, V. Pražák, D. Vasishtan, M. Siggel, F. Hentzschel, A. M. Binder, E. Pietsch, J. Kosinski, F. Frischknecht, T. W. Gilberger, K. Grünewald, Variable microtubule architecture in the malaria parasite, Nat. Commun. 14 (1) (2023) 1216. doi:10.1038/s41467-023-36627-5.

[32] S. Lembo, L. Strauss, D. Cheng, J. Vermeil, M. Siggel, W. C. D. Cheng, J. Vermeil, M. Siggel, M. Toro-Nahuelpan, C. J. Chan, J. Kosinski, M. Piel, O. Du Roure, Others, The distance between the plasma membrane and the actomyosin cortex acts as a nanogate to control cell surface mechanics, bioRxiv (2023) 2001–2023doi:10.1101/2023.01.31.526409.

[33] F. Förster, O. Medalia, N. Zauberman, W. Baumeister, D. Fass, Retrovirus envelope protein complex structure in situ studied by cryoelectron tomography, Proc. Natl. Acad. Sci. 102 (13) (2005) 4729–4734. doi:10.1073/pnas.0409178102.

[34] Napari contributors, napari: a multi-dimensional image viewer for python, doi:10.5281/zenodo.3555620 (2023).

