## Supplementary material for "ColabSeg: An interactive tool for editing, processing, and visualizing membrane segmentations from cryo-ET data": User Guide

### - Supporting Information and User Guide - ColabSeg: An interactive tool for editing, processing, and visualizing membrane segmentations from cryo-ET data

Marc Siggel[a,b], Rasmus K. Jensen[c], Julia Mahamid[c], Jan Kosinski[a,b,c]

[a] *European Molecular Biology Laboratory (EMBL) Hamburg, Notkestrasse 85, Hamburg, 20607, Germany*
[b] *Centre of Structural Systems Biology (CSSB), Notkestrasse 85, Hamburg, 20607, Germany*
[c] *Structural and Computational Biology Unit, European Molecular Biology Laboratory (EMBL) Heidelberg, Meyerhofstrasse, Heidelberg, 60120, Germany*

---

---

#### 1. Setup and Installation

##### *1.1. Setting up ColabSeg*

The Installation is also explained in detail in the Gitlab folder available at: `https://github.com/KosinskiLab/colabseg`. Make an environment with anaconda and open it:

```
conda create --name YOUR_ENV_NAME python==3.8 pip
source activate YOUR_ENV_NAME
```

Run pip in the folder where setup.py is located. This also installs all necessary dependencies:

```
pip install .
```

Then add this environment as jupyter kernel and then boot a new jupyter notebook or better the demo notebook colabseg_demo_notebook.ipynb in the colabseg folder:

```
python -m ipykernel install --user --name=YOUR_ENV_NAME
jupyter notebook colabseg_demo_notebook.ipynb
```

Make sure to pick the correct environment as kernel to have access to the installed software. Execute the cells in order. It is possible to skip the tensorvoting step if segmented data is already available. Then simply load the .mrc file. Alternatively, you can load a .h5 file which is a specific state file of the software which contains all the metadata of the classes. When loading a new file it is advised to either restart the kernel and start from the top of the notebook, or at least re-run the file-loading cell. This will purge any existing data and avoid potential issues in the experimental stage.

##### *1.2. Setting up TomoSegMemTV*

The tool relies heavily on the TomoSegMemTV tool, which is used at the beginning of the pipeline. TomoSegMemTV can be downloaded as an executable directly from the developer's page at this link:
https://sites.google.com/site/3demimageprocessing/tomosegmemtv
(please appropriately cite this work if you use ColabSeg). Note that the path to your downloaded executable of TomoSegMemTV has to be added manually in the GUI's text field (Fig. 1(1)).

#### 2. Features and Sample Usage

##### 2.1. Segmentation with TomoSegMemTV

###### 2.1.1. Usage

Look at Figure 1 for a visual guide. The inputs for ColabSeg can either be prepared with custom scripts using TomoSegMemTV or using the tensor voting GUI provided in the notebook. The optimized settings provided with the GUI are optimized for a px size of approx. 13 Å± 2 Å(But also work very well for up to 26 Å). Other sizes will require different settings since TomoSegMemTV works on a per-voxel basis and the varying pixel size will change how many pixels the relevant membrane features occupy. So far these have proven to work well for a broad range of data sets but need to be adapted on a case-by-case basis. In some cases, it is helpful to pre-filter the data with some denoising tool. The following parameters of the pipeline can be controlled directly in the GUI:

- base_name: The base name of the input image file.
- tensor_voting_path: The path to the tensor voting code.
- cpus: The number of CPUs to use to run the tensor voting. the code can run in parallel. For maximum speed run with all cores of the machine.
- scale_space: The scale space parameter for the tensor voting algorithm used in the gaussian filtering step
- tv1_value: determines the scale factor, i.e., the number of voxels considered in the tensor voting process.
- m_gaussian_pre: The pre-gaussian parameter for the surfaceness calculation.
- m_gaussian_post: The post-gaussian parameter for the surfaceness calculation.
- m_thresh: The threshold value for the surfaceness calculation.
- tv2_value: determines the scale factor, i.e., the number of voxels considered in the tensor voting process.
- s_gaussian_pre: The pre-gaussian parameter for the saliency calculation.

- s_gaussian_post: The post-gaussian parameter for the saliency calculation.
- thresholding_threshhold: The threshold value for the thresholding step.
- cluster_cutoff: The cutoff value for the clustering step.
- remove_intermediates: A boolean parameter indicating whether to remove intermediate files after the segmentation is complete.

The GUI chains together all individual executables of TomoSegMemTV and provides a gzip compressed output file in the output directory. A detailed explanation is available in the TomoSegMemTV paper and user guide (`https://sites.google.com/site/3demimageprocessing/tomosegmemtv`). Users can also use the provided bash script (`https://github.com/KosinskiLab/colabseg`) to run the TomoSegMemTV pipeline.

*2.1.2. Optimized Parameters and Recommendations*

We tested a variety of settings with TomoSegMemTV because some of the defaults did not produce consistent results in particular when scaling to larger data sets. For most of the data sets we used, we recommend a specific set of parameters that produce fairly consistent results and provide a good starting point for downstream optimization. We opt for settings where more features are captured and filter them using the ColabSeg GUI.

```
scale_space -s 2 -f ${in}.mrc ${in}_sspace.mrc
dtvoting -s 15 ${in}_sspace.mrc ${base_name}_tv1.mrc
surfaceness -s 0.8 -p 0.5 -m 0.02 ${in}_tv1.mrc ${in}_surf1.
    ↪ mrc
dtvoting -w -s 10 ${in}_surf1.mrc ${in}_tv2.mrc
surfaceness -S -s 0.75 -p 0.5 -l 10 ${in}_tv2.mrc ${in}_surf2.
    ↪ mrc
thresholding -l 0.06 -2 6 ${in}_surf2.mrc ${in}_thresh.mrc
global_analysis -v 2 -3 3000 ${in}_thresh.mrc ${in}_global2.
    ↪ mrc
```

13-26 Å pixel size work well with these settings and were used in our test case with minor adaptions. The small pixel size also drastically speeds up all steps of the processing including any usage with the ColabSeg GUI. Using -t command it is possible to assign N processes to the task i.e. using multiple cores. The GUI is automatically populated with the optimized values and can be directly run by the user.

##### 2.2. Point cloud conversion

Now look at figure 2. The output the TomoSegMemTV wrapper provided in the notebook or the final output of the bash scripts is a .mrc file that only contains integer values i.e. 0 where no membrane present and integer $\{1..N\}$ for each membrane cluster. Any other mrc file following this convention can be imported. In case another file was generated with any other segmentation tool e.g. using a machine learning output using the TomoSegMemTV connected_component executable on the file can be useful to obtain the correct file format and pre-cluster all the pieces of the file. This also ensures that any potential issues with the file format are fixed. The file (relative or absolute path) can be designated in the GUI and uploaded into the point cloud converter. The conversion generates xyz coordinates from the mrc file and can then be loaded in the following step into the GUI. Note that large numbers of clusters $> 100$ can take a very long time to process. In that case, it is also possible to run this on a cluster by directly accessing the conversion function "convert_tomo". After this loading step, the system is ready to be manipulated and processed in the ColabSeg GUI.

##### 2.3. Viewer and cluster management

Now check figure 3 for a visual guide. With the data prepared the viewer can be loaded in the next step and the segmentation visualized with the point cloud viewer. The viewer automatically downsamples the number of shown voxels if the number becomes prohibitively large to avoid crashing the browser. If the points seem far apart the reason is this downsampling procedure. However, when writing the full file this is not an issue because the full dataset is considered. The cluster window gives a numbered list of clusters that are initially ordered by size. Note however that when deleting clusters or merging them this order will be changed. When the cluster is selected in the list viewer it is highlighted in red.

It is possible to select multiple clusters by clicking the command button and selecting multiple entries. By clicking merge clusters it is possible to merge these clusters into one group. Any amount of clusters can be merged. The clusters are then renumbered. This is important later if membranes of different types are present in the data. In our example, we need to merge for instance as an extended piece of membrane is patchy and multiple clusters define the same membrane. One can merge them in this way for further processing with fits. Similarly, clusters can be deleted if they aren't of interest. Removing uninteresting clusters first is helpful as this makes ColabSeg run faster.

*2.4. Undoing step and reloading initial data*

Figure 3 provides a visual guide for undoing steps or reloading data. If during the edition process, a step was performed by accident or the results are unsatisfactory, it is possible to undo this by clicking the undo button on the cluster management page. This restores the previous step. Note that this is only possible ONCE! Older steps are not saved. To be sure not to lose your progress along the way, use the hdf5 file to save the state of the entire GUI to a file.

Alternatively, it is possible to restore the original state after loading and converting the tomogram using the restore initial button. This is particularly useful if multiple editing steps have been performed.

*2.5. Lamella editing*

Lamella editing tab is shown in Figure 6. This method takes a list of cluster indices and trims the top and bottom of each cluster along a given axis. The amount to trim is specified by trim_min and trim_max parameters, which are added and subtracted from the minimum and maximum value of the given axis, respectively. The axis to trim along is specified by the trim_axis parameter, which can take values of ”x”, ”y”, or ”z”. If an invalid value is provided, an exception is raised. For each cluster in the provided list of indices, the method first calculates the minimum and maximum values along the specified trim axis. This new, trimmed array replaces the original cluster.

*2.6. Filtering and processing clusters*

Now we move on to Figure 5. The output from TomoSegMemTV often has issues from artifacts, such as fiducials or other contaminations giving false positives which have to be filtered. Similarly, membranes on occasion are very close together and are not properly separated with TomoSegMemTV on its own. Possibly settings exist which enable filtering these issues but usually, this would need to be tuned for each tomogram. Therefore, features for filtering are provided to the user which can also be used for batch processing. In any case a majority of the manual labor of manual cleaning with software such as Amira should be alleviated with these features. Our test show that this works quite well for many cases, especially vesicles, viruses, and long planar membranes such as the plasma membrane. The processing tab can be opened and all these features are available there. To process a cluster first the cluster of interest needs to be selected in the cluster list. The procedures only work on a single cluster.

1. statistical_outlier_removal: This function removes statistical outliers from a cluster of points. It takes in a cluster_index parameter that specifies which cluster to process, a nb_neighbors parameter that specifies the number of neighbors to consider when computing the mean distance, and a std_ratio parameter that specifies the standard deviation multiplier for distance. The function uses the Open3D library's remove_statistical_outlier function to perform the outlier removal and updates the cluster with the remaining points.
2. dbscan_clustering: This function performs DBSCAN clustering on a cluster of points. It takes in a cluster_index parameter that specifies which cluster to process, a 'minimal_dbscan_size' parameter that specifies the minimum number of points required for a subcluster to be created, an eps parameter that specifies the maximum distance between two points to be considered in the same neighborhood, and a min_points parameter that specifies the minimum number of points required to form a dense region. The function uses the Open3D library's cluster_dbscan function to perform the clustering and updates the cluster with the resulting subclusters.
3. outlier_removal: This function takes in a cluster index, a number of nearest neighbors ($k_n$), and a threshold value as inputs. It uses covariance-based edge detection to remove points from the point cloud. The function first converts the input point cloud to a PyntCloud object, calculates the eigenvalues and eigenvectors for each point, and then uses the eigenvalues to detect edges. Points with eigenvalues below the specified threshold are removed from the point cloud. The resulting point cloud is then saved and returned.

A workflow that has worked well is first trimming possible fuzzy edges of the z-stack, which might lead to the merging of adjacent membranes. In case the tilt was not corrected during the reconstruction process, this can be done here in postprocessing by using the rotation function. After trimming it can also be rotated back to its original angle such that the segmentation fits with the raw data in case the outputs need to be overlaid with other data. Next, it is helpful to run the edge outlier removal to remove possible features which are highly irregular or have a fast-changing curvature field. In principle, some parameters can be adjusted and optimized for each tomogram. However, after rigorous testing across numerous tomograms, we found the settings worked best with the presets provided. Therefore, no settings can be changed here. Ideally, points that connect the clusters which were falsely connected are removed. Then it is possible to recluster these with the DBSCAN method. If successful, the number of total clusters will increase drastically. The parameter defaults for DBSCAN work well with a $\approx$ 13 Åpixel size (also works for up to 26 Å). If the pixel size is larger the radius parameter and the minimal points parameter need to be increased to get a sufficient coverage and not lose too many points. This workflow usually works quite well. If the edge outlier removal is not sufficient, it is also possible to use the statistical outlier removal strategy. This is often more aggressive and removes more points but enables proper membrane separation. The smaller the sigma parameter is chosen the more points are removed from the segmentation. The parameters can be varied. Here it has proven helpful to test a parameter and if necessary undoing the step by returning to the cluster selection menu and clicking the undo button.

##### *2.7. Fitting Membranes*

The visual guide for fitting can be found in Figure 6. ColabSeg allows fitting clusters with radial basis function (RBF) fits if the membranes are very planar or spheres for vesicles or viruses. To use this feature first select the cluster you want to fit from the cluster list. Click on the fitting tab next and choose the fitting method, either for a sphere or using the RBF fit. Important for the RBF is to choose the correct plane in which the membrane lies. The fit chooses the coordinate system accordingly to apply the fit properly. In some instances, the fit is off at primarily the edges and this can lead to issues when fitting the data. Therefore, it is also possible to fix this by removing all points which are further from the original segmentation than a defined distance. This can be helpful to trim possible inaccuracies which can be introduced through the RBF fit. Note that the max distance should be chosen that any holes in the membranes are properly patched.

##### *2.8. Saving/Exporting Files*

Saving and exporting is shown in Figure 7. When all the processing is done the files can be saved as various file output formats. The file formats are a text file that saves the positions as XYZ coordinates. Alternatively, the data can be written as an mrc file for further processing. Specify a filename in the output filename text box. By default, ColabSeg only writes those clusters and fits which were selected manually. If further analysis is only to be performed on a subset of the membranes it is possible to save individual or merged clusters.

##### *2.9. Saving a Session*

If you want to continue editing and want to save the session for further edits at a later point it is possible to save a session using an hdf5 file format which dumps the state of the backend into this file. The file ending should be ".h5". This file format can be reloaded in the conversion section above as an alternative entry point to populate the GUI with data.

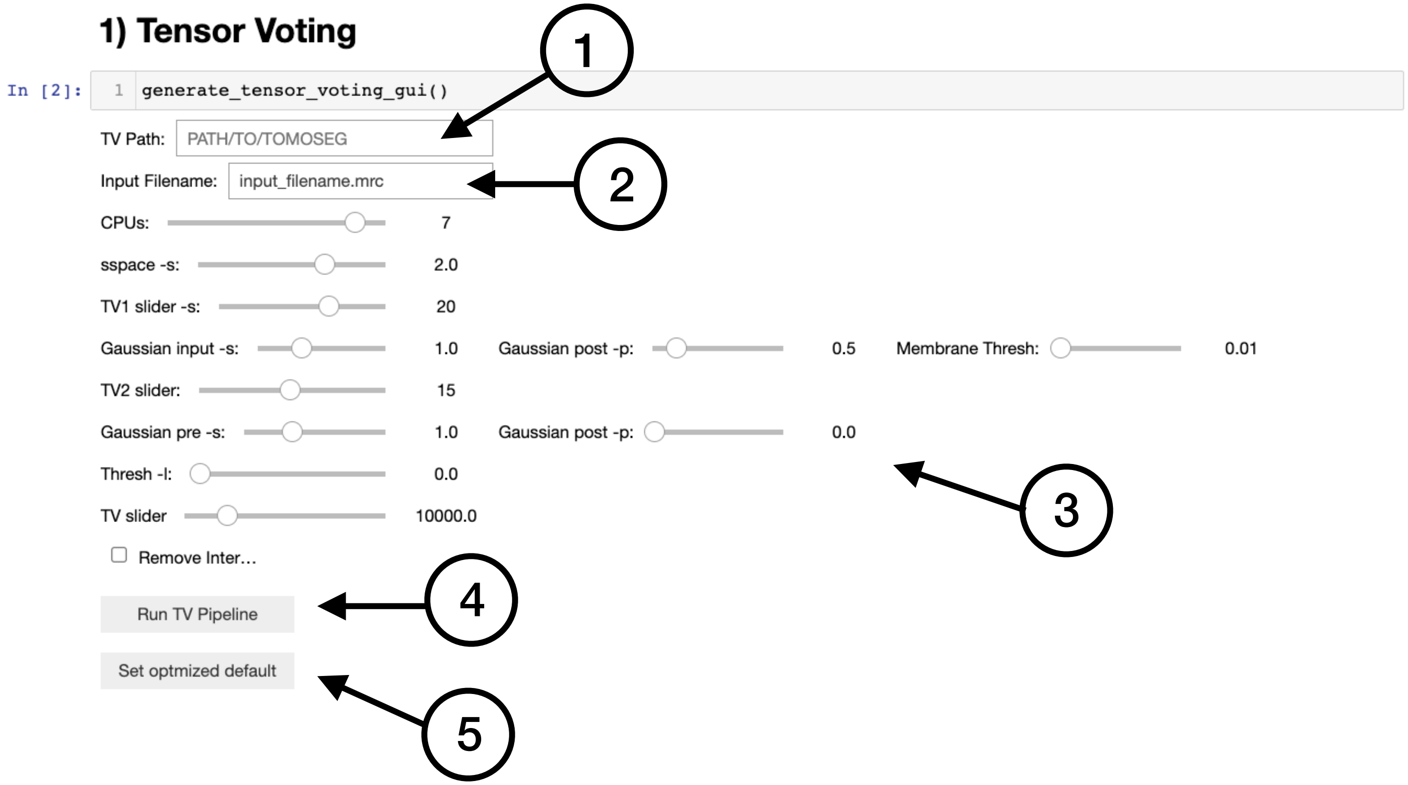

Figure 1: TensorVoting Menu. (**1**) Field to provide the absolute path to the folder containing the TomoSegMemTV executables. All must be located in the same folder, and the names kept as downloaded. (**2**) Input for the relative or absolute path of the input .mrc file to be processed in the pipeline. (**3**) Settings according to the manual of TomoSegMemTV. The slides are ordered in the sequence of the steps as they are applied as recommended by TomoSegMemTV. (**4**) Runs the complete tensor voting pipeline with the settings provided. (**5**) Sets all sliders to optimized settings which work well for a 13 Å/voxel size .

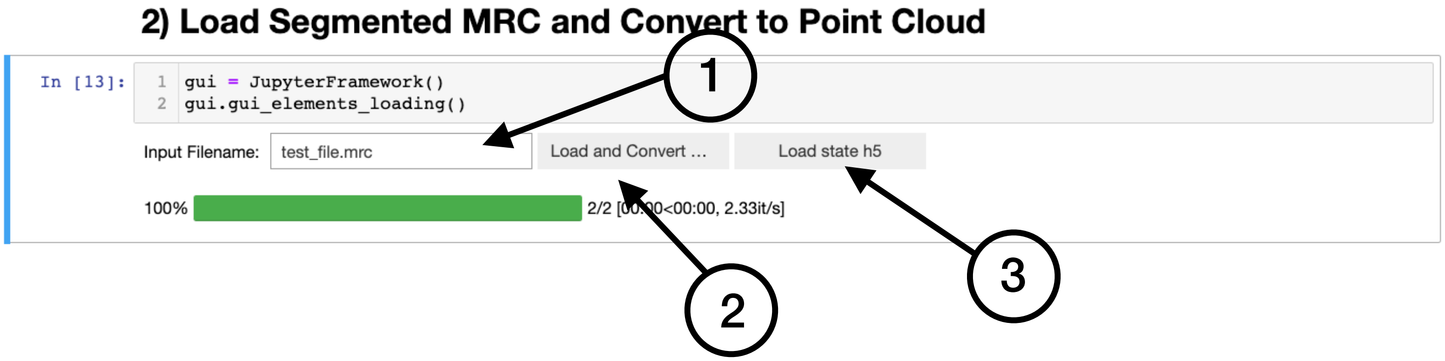

Figure 2: Data Loading Menu. (**1**) Relative or absolute path for the file to be loaded in the UI. (**2**) The status bar indicates the time to process the tomogram. Large mrc files with many clusters and many voxels can take a long time to process (time indicated). We recommend working with a larger pixel size and upsampling later. (**3**) Alternative to the segmentation it is possible to load a state file from a previous session. Only hdf5 files generated with ColabSeg (see saving) can be loaded here. This restores the complete state of the GUI.

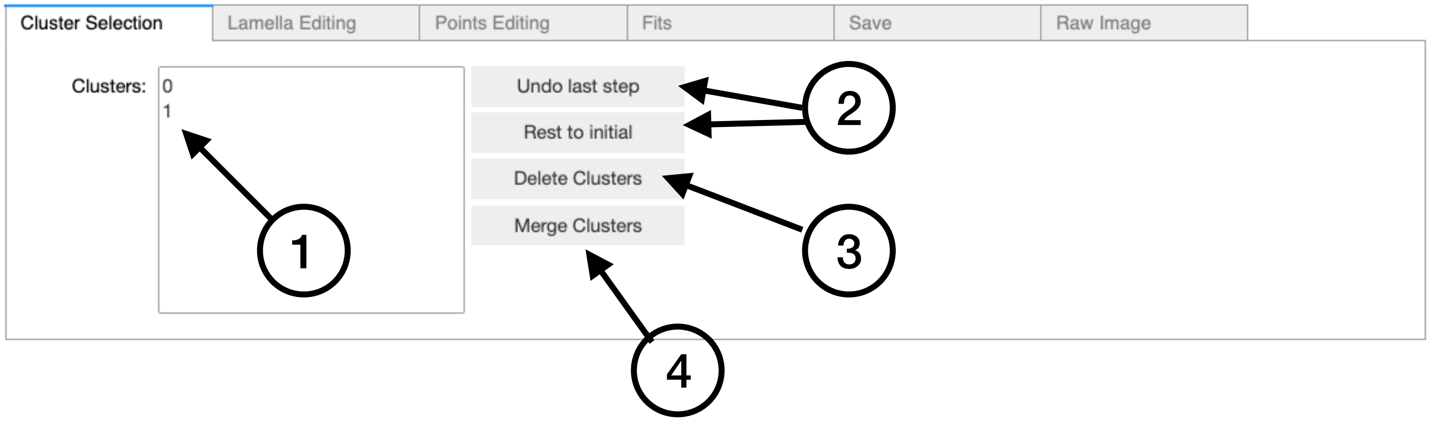

Figure 3: Cluster Selection Tab. **1** Listing to select clusters. Multi-select is possible. the clusters are highlighted in red in the 3D viewer if they are selected. **2** Tools to undo the last step or reload the initial state of the GUI, which can undo all previous processing steps. **3** Delete one or multiple clusters which are selected in the cluster selection window (1).**4** Merge all selected clusters into one. Note that the numbering of the clusters can change.

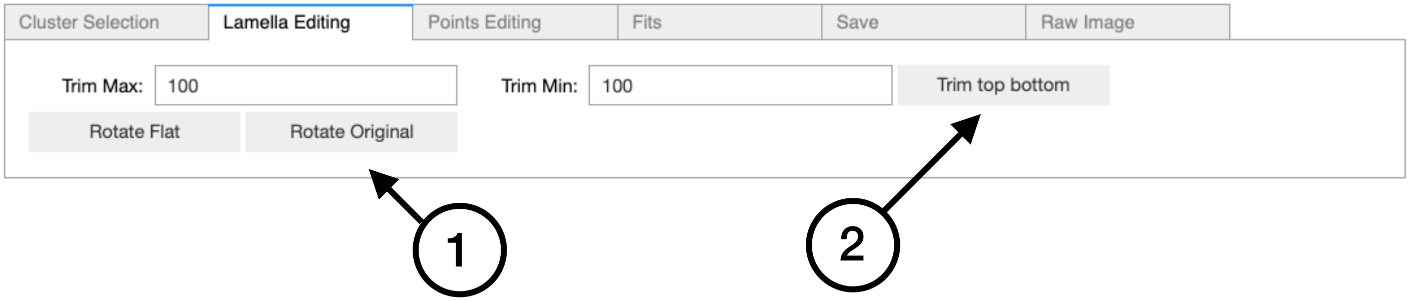

Figure 4: Lamella Editing Tab. **1** Aligns the global plane of the lamella with the xy plane if they were not corrected during reconstruction. This is useful for trimming. The rotation can be restored to the original angle to overlay with raw data. **2** Trimming of the selected clusters at the top and bottom according to the values listed in trim max and trim min. The trimming is relative to the highest and lowest point of the current height of the voxels. i.e. if applied multiple times, the z thickness of the selected membrane clusters is continuously decreased.

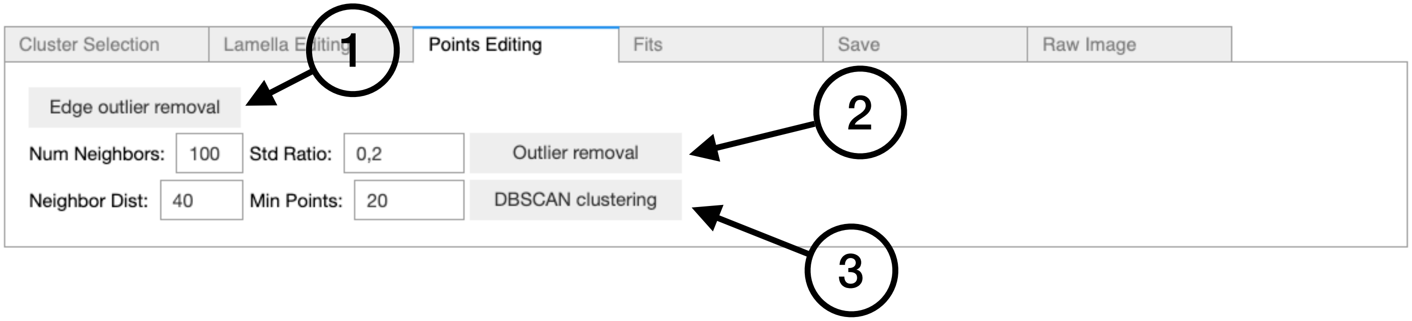

Figure 5: Points Editing Tab. (**1**) Edge outlier removal tool. Runs directly on the selected cluster. The parameters for this tool were optimized internally. Thus, they are not visible here. (**2**) Statistical outlier removal tool. Options are the number of neighbors considered when calculating the average distribution around a point. The smaller the std. ratio and smaller the neighborhood the more selective the filter becomes. (**3**) DBSCAN reclustering allows a single selected cluster to be reclustered. Useful after processing or when using input data that hasn't been checked with a connected component algorithm. Neighborhood distance i.e. radius around each considered point can be set along with the minimal amount of points to classify a point as part of the

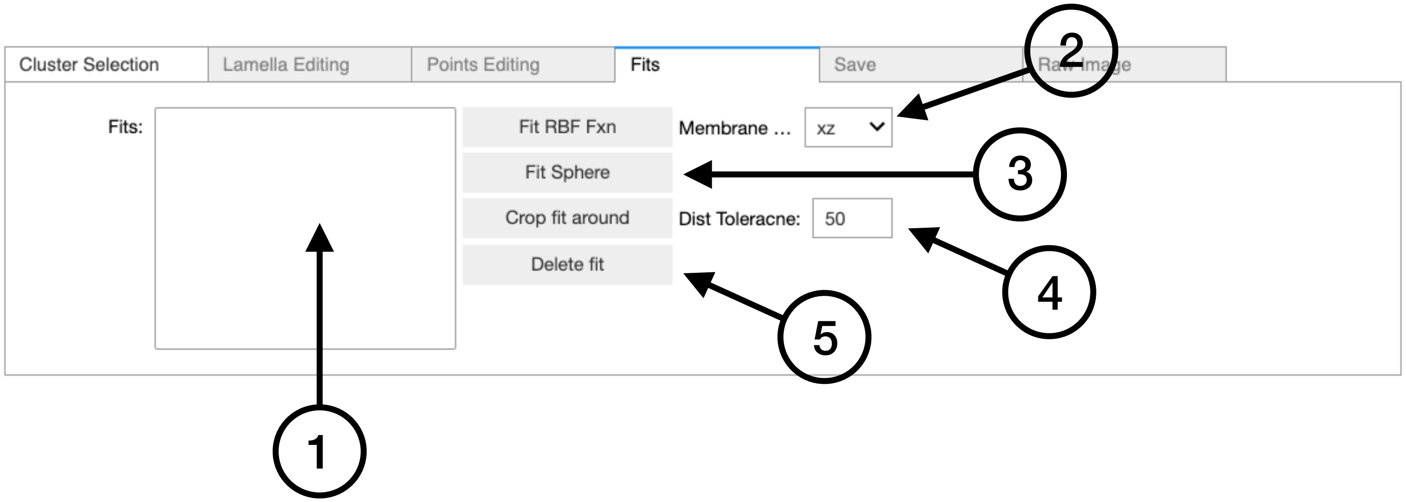

Figure 6: Fitting Tab. (**1**) Window showing all fits as clusters. As in the cluster selector individual fits can be picked and modulated. (**2**) RBF for near-planar membranes. The place orientation gives the direction primary expanse of the plain. This needs to be adapted otherwise the fit won't be correct. (**3**) Sphere fit for a cluster. (**4**) Crops the fit around the initial cluster. This is useful if, e.g., anything beyond the edges of the lamella shouldn't be considered or the extrapolation is too extreme. The distance tolerance sets the distance to the original cluster to which the fit was applied (unit is Å). (**5**) Possibility to delete one or multiple fits from the list if they are no longer needed.

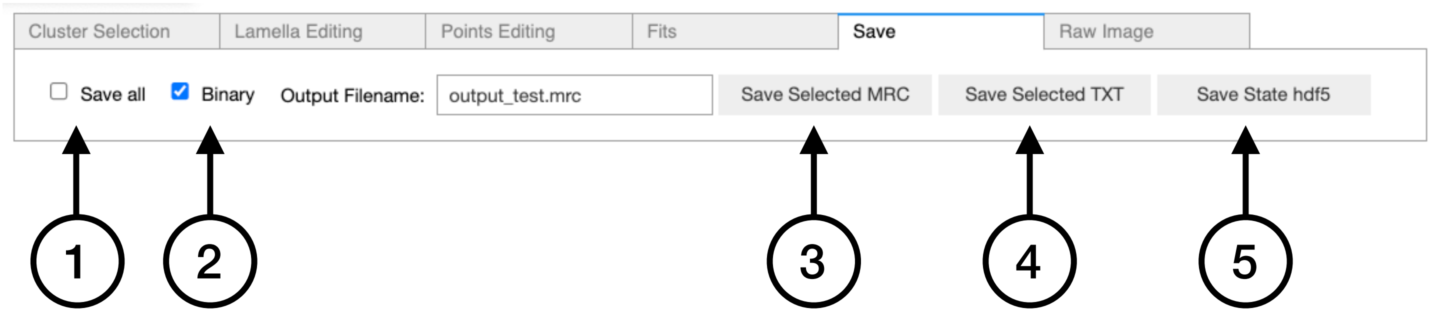

Figure 7: Saving Tab. **1** Checkbox if selected saves all clusters. if not only the ones selected in the GUI. **2** outputs only 0 and 1 for all clusters and ignores the integers assigned to the clusters (many machine learning applications need this format for proper usage). **3** Save the selected files as .mrc file. **4** Save the point coordinates in an xyz file (useful output to use for geometric analyses in real space). **5** Save the state as hdf5 state file. This saves all metadata of the GUI and can only be loaded reasonably within the GUI (useful if more editing is needed at a later time point).
